# PQM-1 controls hypoxic survival via regulation of lipid metabolism

**DOI:** 10.1101/769927

**Authors:** Thomas Heimbucher, Julian Hog, Coleen T. Murphy

## Abstract

Animals have evolved responses to low oxygen conditions to ensure their survival. Here we have identified the *C. elegans* zinc finger transcription factor PQM-1 as a regulator of the hypoxic stress response. PQM-1 is required for the longevity of insulin signaling mutants, but surprisingly, loss of PQM-1 increases survival under hypoxic conditions. PQM-1 functions as a metabolic regulator by controlling oxygen consumption rates, suppressing hypoxic glycogen levels, and inhibiting the expression of the sorbitol dehydrogenase-1 SODH-1, a crucial enzyme for sugar metabolism. PQM-1 promotes intestinal fat metabolism by activating the expression of the stearoyl-CoA desaturase FAT-7, an oxygen consuming, rate-limiting enzyme in fatty acid biosynthesis. PQM-1 activity enhances fat accumulation in embryos under hypoxic conditions, thereby increasing survival rates of arrested progeny during hypoxia. Thus, while *pqm-1* mutants increase survival of mothers, ultimately this loss is detrimental to progeny survival. Our data support a model in which PQM-1 controls a trade-off between lipid metabolic activity in the mother and her progeny to promote the survival of the species under hypoxic conditions.

## Introduction

A constant oxygen supply is essential to sustain the life of aerobic organisms, which have evolved multiple adaptive mechanisms to maintain the delicate balance between oxygen supply and demand. Impairment of this balance is associated with many age-related diseases, affecting pulmonary and cardiac function^1^ and is increasing the economic cost of aging populations in modern societies. Consequently, understanding the defense mechanisms that animals have developed to protect against oxygen deprivation is crucial for the development of treatment strategies for human ischemia-related diseases and cancer.

*C. elegans* has been used as model to study survival under a range of oxygen levels^2^ because it is well adapted to limited oxygen conditions; in fact, worms prefer 5-12% rather than atmospheric (21%) levels of oxygen^3,4^. In its natural habitat, *C. elegans* frequently encounters conditions where oxygen is depleted, because flooded soils can become hypoxic^5,6^. When exposed to very low oxygen levels (<0.4% O_2_), *C. elegans* enters a state of suspended animation^2^, a reversible hypometabolic state in which biological processes are largely slowed or even arrested. Intermediate levels of oxygen differentially affect transcriptional responses. The key hypoxic transcriptional regulator HIF-1 is required for development and survival of *C. elegans* at 0.5% - 1% oxygen (hypoxia), whereas it is not essential for survival in complete anoxia^7^. More recently, the zinc finger protein BLMP-1 was identified as a regulator of a HIF-1-independent hypoxic response based on a reporter screen performed with the hypoxia mimetic CoCl_2_, which largely replicates the hypoxic state^8^.

Insulin/IGF-1 signaling (IIS) controls survival under various stresses, including hypoxia. Reduction-of-function mutations in the Insulin/IGF-1 receptor homolog *daf-2* protect *C. elegans* from high-temperature hypoxia and long-term anoxia^9,10^. The resistance against oxygen depletion is mediated by the FoxO transcription factor DAF-16, the major downstream transcriptional effector of IIS^11–13^; ^14,15^.

We previously identified the C2H2-type zinc finger protein PQM-1 as an IIS-regulated DAF-16 antagonist and transcriptional regulator that is partially required for the exceptional longevity of *daf-2* mutants^15^. As reduction of IIS is highly protective for survival of nematodes exposed to hypoxic stress^9^, here we asked whether PQM-1, like DAF-2 and DAF-16, plays a role in hypoxic survival. Surprisingly, we found that loss of *pqm-1*, unlike loss of *daf-16*, protects *C. elegans* from hypoxic stress. To understand the underlying mechanisms of this protection, we studied the transcriptional changes in *pqm-1* mutants under normoxia versus hypoxia and discovered that alterations in carbohydrate and lipid metabolism are key to this survival. Ultimately, however, the loss of *pqm-1* under hypoxic stress is detrimental to future generations, suggesting that PQM-1 is an important component of multi-generational hypoxic survival through its regulation of key lipid metabolic genes.

## Results

### PQM-1 acts as a negative regulator of hypoxic survival independently of Insulin-like signaling

We previously found that the zinc finger transcription factor PQM-1 is required for *daf-2* mutants’ extended longevity^15^, and thus we expected PQM-1 to also be required for *daf-2’s* extended survival in hypoxic conditions. Exposure of *C. elegans* to the hypoxia mimetic cobalt chloride (CoCl_2_) activates HIF-1 and mimics hypoxic responses^16^, allowing timecourse survival analysis under hypoxia-like conditions^8^, whereas a typical hypoxic chamber experiment allows one survival time point after a normoxic recovery period. Counter to our expectations, *pqm-1(ok485);daf-2(e1370)* double mutants exposed to 5 mM CoCl_2_ survived significantly longer than did *daf-2(e1370)* worms (Fig. 1a). Similarly, knockdown of *pqm-1* via RNA interference further increased the hypoxia survival of *daf-2(e1370)* mutants (Fig. 1b). Together, these results suggest that PQM-1 activity may limit resistance of *daf-2(e1370)* animals exposed to hypoxia-like stress.

**Figure 1.**
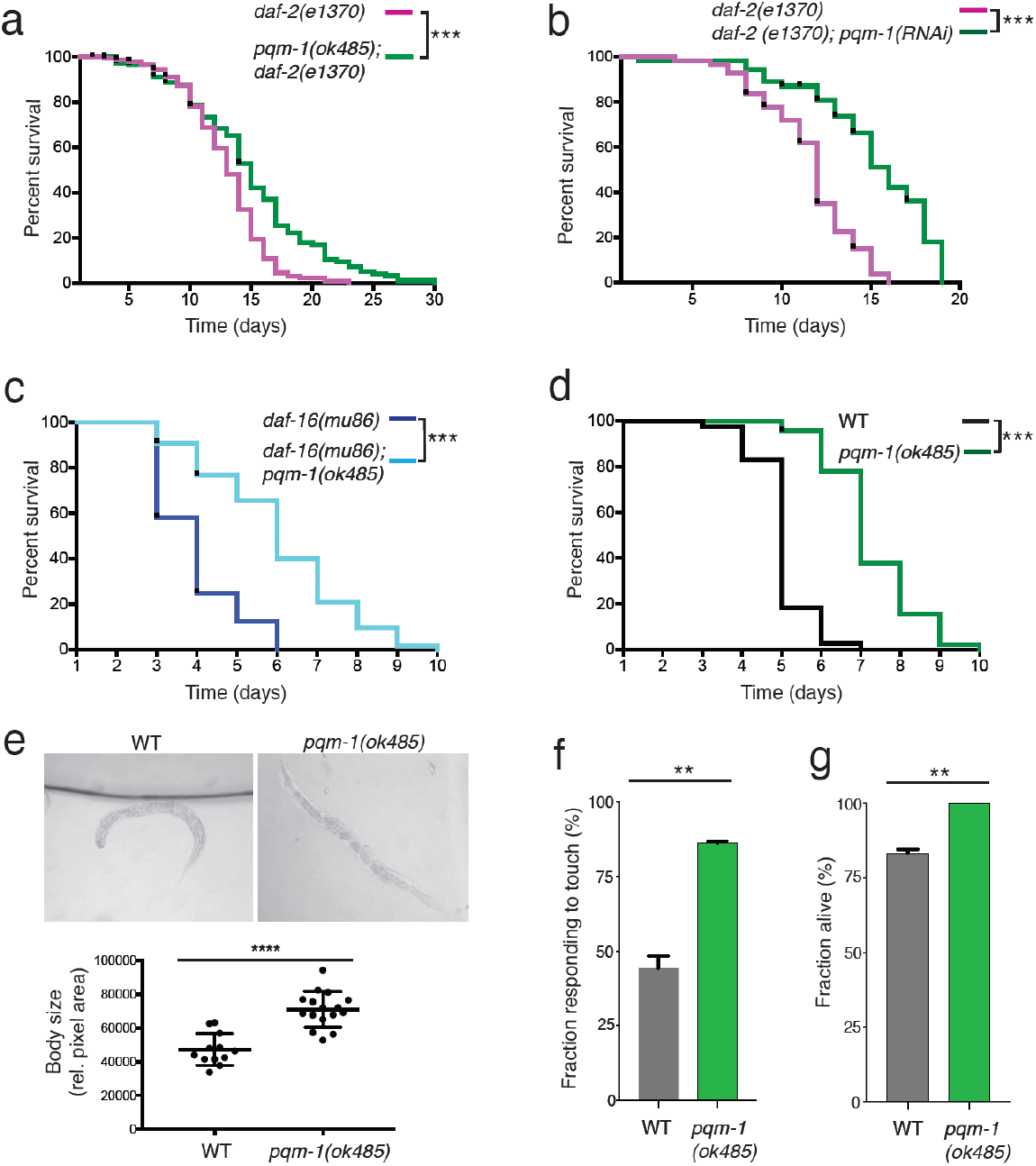
Loss of *pqm-1* increases survival and recovery under hypoxic conditions independently of IIS and FOXO. a-d) Survival analysis of animals exposed to 5 mM CoCl_2._ 45 ≤ n ≤ 78, * p<0.02, *** p<0.0001, Log-Rank analysis was performed. Three independent experiments were performed. e) Shrinking of worms exposed to 5 mM CoCl_2_ for 3 days and quantification of body size. f) Measurement of touch response of animals after exposure to 0.3% O2 in a hypoxic chamber for 24 hr at room temperature followed by a 30-minute recovery period. g) Survival analysis after exposure of animals to 0.3% O_2_ in a hypoxic chamber for 25 hours at 26°C followed by a one-day recovery period. e-g) Two-tailed t-test, mean ± SEM, n ≥ 37, ** p<0.005, **** p<0.0001.

The transcriptional outputs of IIS are largely mediated by the FOXO transcription factor DAF-16^14,15^. *daf-16* and *pqm-1* act similarly in longevity regulation, as the mutants both have normal/short lifespans and both transcription factors are required for *daf-2’s* longevity. However, under hypoxia-like stress, inactivation of *pqm-1* greatly increased the survival of a *daf-16(mu86)* null mutant (p<0.0001; Fig. 1c and S1), suggesting that PQM-1 acts independently of DAF-16 to regulate hypoxic survival. *pqm-1(ok485)* loss-of-function mutants also survived longer than wild-type animals (Fig. 1d and S1) and maintained a largely normal morphology, whereas wild type worms shrank after 3 days of exposure to the hypoxia mimetic (Fig. 1e). Our data indicate that PQM-1 acts as a negative regulator of somatic integrity and survival under hypoxic stress.

To verify our findings from CoCl_2_-induced hypoxia-like conditions to true hypoxia, we tested the recovery rate of wild-type and *pqm-1(ok485)* mutants after treatment in a hypoxic chamber. When animals were exposed to 0.3% O_2_ for 24 hours at room temperature followed by a short normoxic recovery period, *pqm-1(ok485)* mutants exited suspended animation earlier than did wild type (Fig. 1f). Similarly, *pqm-1(ok485)* mutants survive oxygen depletion better than do wild-type animals (Fig. 1g) upon reduction of oxygen levels combined with increased temperature^9^. Together, these results suggest that loss of PQM-1 function improves survival under both hypoxia and CoCl_2_ hypoxia-like conditions.

### PQM-1 is a transcriptional regulator of the hypoxic stress response

Under normoxia, the transcription factor PQM-1 controls gene expression programs that regulate development, growth, and reproduction^15^. To identify genes altered in hypoxia-like conditions, we carried out two types of analyses: first, a direct comparison between *pqm-1* and wild-type animals that had both been exposed to CoCl_2_ to identify those genes that provide *pqm-1* a survival advantage under hypoxic conditions (Fig. 2a, Table S2; One-class SAM, 1650 upregulated and 629 downregulated genes). Secondly, we carried out a Two-class SAM analysis in comparisons of wild-type and *pqm-1* worms both treated with CoCl_2_ vs untreated to further identify potential *pqm-1* targets that increase survival under hypoxic conditions (Fig. 2b, c; Table S3). We found that genes associated with heat shock response (*hsp-16*, *hsp-70*), metal response (*numr* and *cdr*), infection response genes (*irg-1*, *-2*), glutathione S-transferases (*gst*), several DAF-16 target genes (*dod-22, -17, -24*), and PQM-1/DAF-16 regulated targets (C32H11, K12H11, and F55G11)^14^ are upregulated, indicating that the animals are responding to stress conditions. Several cytochrome P450s (*cyp-35A2, -25A1, -14A2, -25A2, -34A9/dod-16*), short chain dehydrogenases (*dhs-25, -20, -2*) and lipid metabolism genes (e.g., *lips-14, elo-6, acdh-2, elo-5*) are downregulated under hypoxia-like conditions (Fig. S2a; Table S1; FDR < 0.01), indicating metabolic changes. Gene Ontology analysis (Fig. S2b) suggests that worms subjected to CoCl_2_ hypoxic conditions affect immune and defense response, ER signaling, aging and lifespan regulation, and lipid metabolism, while downregulated genes are associated with the response to xenobiotic stimuli, oxidation-reduction processes, lipid metabolism, and transmembrane transport (Fig. S2c,d; Table S2). GO analysis of the *pqm-1* and CoCl_2_-dependent upregulated gene sets revealed an enrichment for oxidation-reduction processes and the innate immune response (Fig. S3a). GO term “metabolism” is highly significant in the downregulated gene set (Fig. S3b), with fatty acid metabolism, flavonoid related processes, and transmembrane transport GO terms statistically enriched.

**Figure 2.**
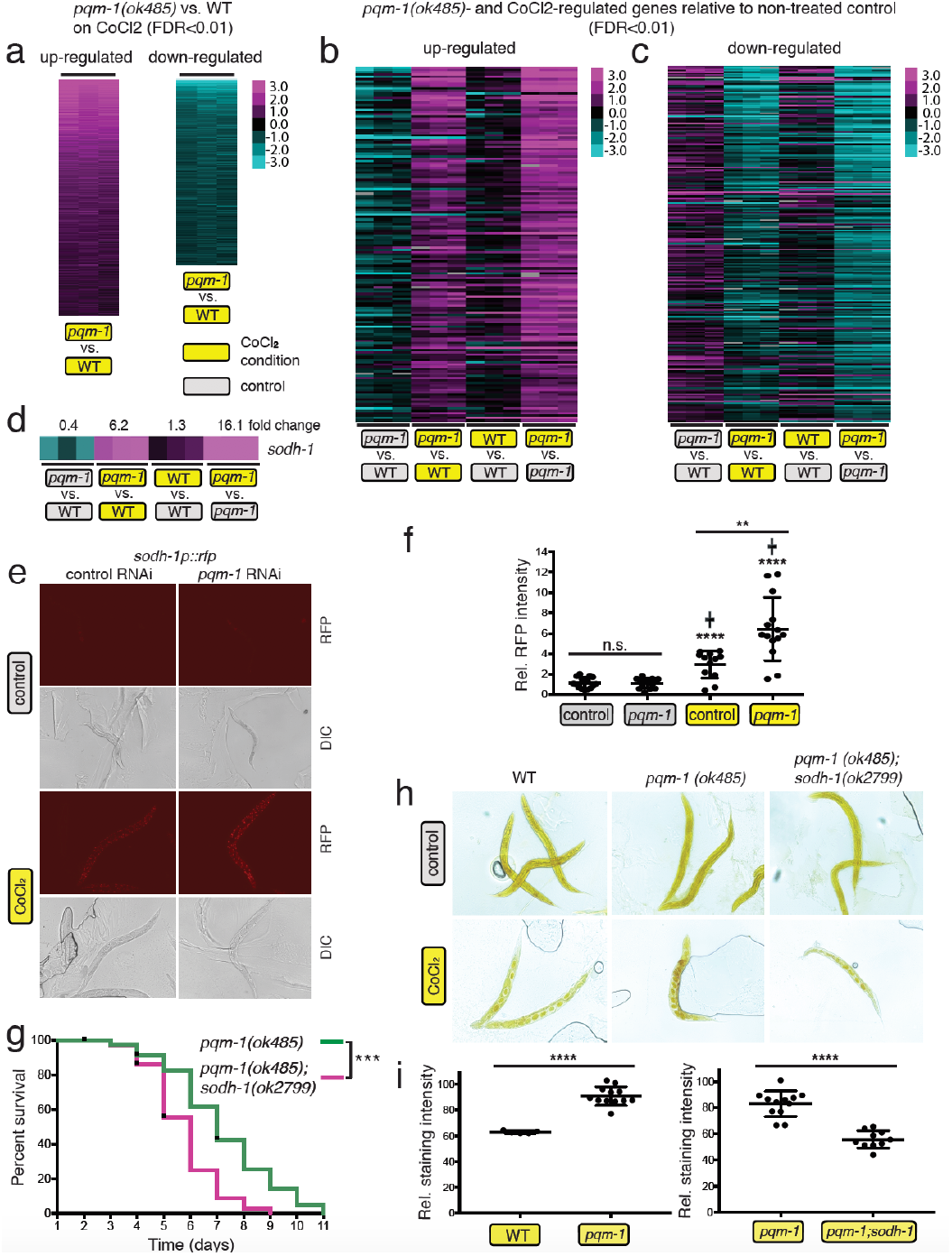
Glycogen metabolism is altered under hypoxia via PQM-1’s regulation of *sodh-1*. a) One-Class SAM analysis of gene expression changes in *pqm-1(lf)* +CoCl_2_ versus WT +CoCl_2_ identified 1650 upregulated and 629 downregulated genes (Table S2). b,c) *pqm-1-* and CoCl_2_- dependent up-regulated (b) and down-regulated genes (c). Two-Class SAM of *pqm-1* +CoCl_2_ versus WT +CoCl_2_ compared to untreated *pqm-1(lf)* versus untreated WT identified 152 upregulated and 243 downregulated genes (Table S3). Heat maps depict individual genes (rows) ranked by SAM score (median FDR < 0.01), with columns representing individual microarrays (a, b, c). Animals were exposed to 5 mM CoCl_2_ for 6 hr at early adulthood (a,b,c). d) *pqm-1-* and CoCl_2_-dependent expression changes for endogenous *sodh-1*. e) *sodh-1::rfp* reporter activity after 20 hr of CoCl_2_ exposure; (f) quantification of reporter activity in (e). Two-tailed *t*-test, mean ± SEM, ** p<0.01, **** p<0.0001, ns = not significant, (+) indicates CoCl_2_-exposed empty vector control and CoCl_2_-exposed *pqm-1* knock down condition versus non-treated empty vector control and non-treated *pqm-1* knock down condition, respectively. g) Survival analysis of *pqm-1* and *pqm-1;sodh-1* mutants exposed to 5 mM CoCl_2_. n ≥ 56, ***p<0.0001 (Log-Rank analysis). Three independent experiments were performed. h,i) Glycogen levels in wild type, *pqm-1*, and *pqm-1;sodh-1* mutants subjected to 5 mM CoCl_2_ for 3 days followed by iodine vapor staining (h); (i) quantification of glycogen levels in (h). Two-tailed *t*-test, mean ± SEM, **** p<0.0001.

Carbohydrate metabolism is critical for *C. elegans* survival under conditions where oxygen is limited^10,17,18^. *sodh-1/dod-11* was one of the top two-class SAM hits for upregulated genes in a *pqm-1(lf)* mutant challenged with CoCl_2_ (Fig. 2d, Table S3). Sorbitol dehydrogenase (SODH) enzymatic activity converts sorbitol, the sugar alcohol form of glucose, into fructose^19^. *sodh-1/dod-11* was previously identified as a downstream target of the IIS/DAF-16 pathway and is required for the long lifespan of *daf-2(lf*) mutants^14^; moreover, it is a key enzyme for starvation-induced aggregation of *C. elegans* and ethanol metabolism^20^ and is upregulated when worms are exposed to transient hypoxia^21^. The promotor region of *sodh-1/dod-11* contains 2 PQM-1 binding sites (DAEs), suggesting that it functions as a direct PQM-1 target gene^14,22^.

We confirmed the upregulation of *sodh-1/dod-11* in a *pqm-1(lf)* mutant upon CoCl_2_ exposure using a fluorescent reporter (Fig. 2e,f), suggesting that PQM-1 likely normally represses *sodh-1* expression. *sodh-1* inactivation in a *pqm-1(lf)* mutant significantly reduced CoCl_2_ survival relative to *pqm-1(lf)* (Fig. 2g, Fig. S3c), indicating that *sodh-1* is necessary for *pqm-1’s* enhanced hypoxic survival. Previous studies have demonstrated that sorbitol, the substrate of SODH-1, can be metabolized to glycogen in diapause eggs of insects (*Bombyx mori*) depending on the activity of SODH-1^23^. To test the role of *pqm-1* and its downstream negatively regulated target *sodh-1* in hypoxic carbohydrate metabolism, we measured glycogen levels. *pqm-1(lf)* mutants maintain glycogen at higher levels than wild-type animals challenged with CoCl_2_ (Fig. 2h,i, Fig. S3d,e), while *pqm-1(lf);sodh-1(lf)* double mutants had reduced glycogen levels, suggesting that *pqm-1’s* increased hypoxia survival is linked to its increased glycogen levels, which in turn depends on SODH-1 function.

### PQM-1 activity regulates lipid levels in hypoxia

The CoCl_2_-induced *pqm-1*-differentially expressed gene list is enriched for lipid metabolism regulators. The downregulated genes included fatty acid elongases (*elo-2, -5, -6*, and -*9*), the stearoyl-CoA fatty acid desaturase *fat-7,* and enzymes functioning in fatty acid β-oxidation (*acdh-1*, *acdh-2, ech-7* and *-9*) (Fig. 3a, top), while upregulated genes included the fatty acid and retinoid binding protein *far-7* and the acyl-CoA synthetases *asc-2* and *asc-17* (Fig. 3a, bottom). Oil Red O lipid staining revealed no significant differences between wild type and *pqm-1* mutants under control conditions (Fig. 3b, c top), but 5 mM CoCl_2_ treatment for 44 hours caused wild-type animals to lose fat relative to the untreated controls (Fig. 3b, d, left). This effect of lipid loss was significantly more pronounced in *pqm-1(ok485)* mutants challenged with CoCl_2_ (Fig. 3b,d right), suggesting that PQM-1 normally acts as a positive regulator of lipid level maintenance. Similarly, *pqm-1* mutants exposed to 0.4% O_2_ (hypoxia) for 44 hours exhibited a significant reduction in fat levels compared to hypoxic wild-type animals (Fig. 3e, f), confirming that CoCl_2_ conditions mimic true hypoxia for fat metabolism. In some individual hypoxic *pqm-1(lf)* animals, lipid levels were completely depleted in the intestinal tissue (Fig. 3e), and fat was only detected in the eggs located in the uterus of *pqm-1(ok485)* mutants. These results suggest that PQM-1 acts as an essential intestinal metabolic regulator, promoting lipid levels under oxygen depletion. However, an additional mechanism might be implicated in maintaining hypoxic fat mass, as wild-type animals also lose fat in a PQM-1-independent manner when challenged with the hypoxia mimetic.

**Figure 3.**
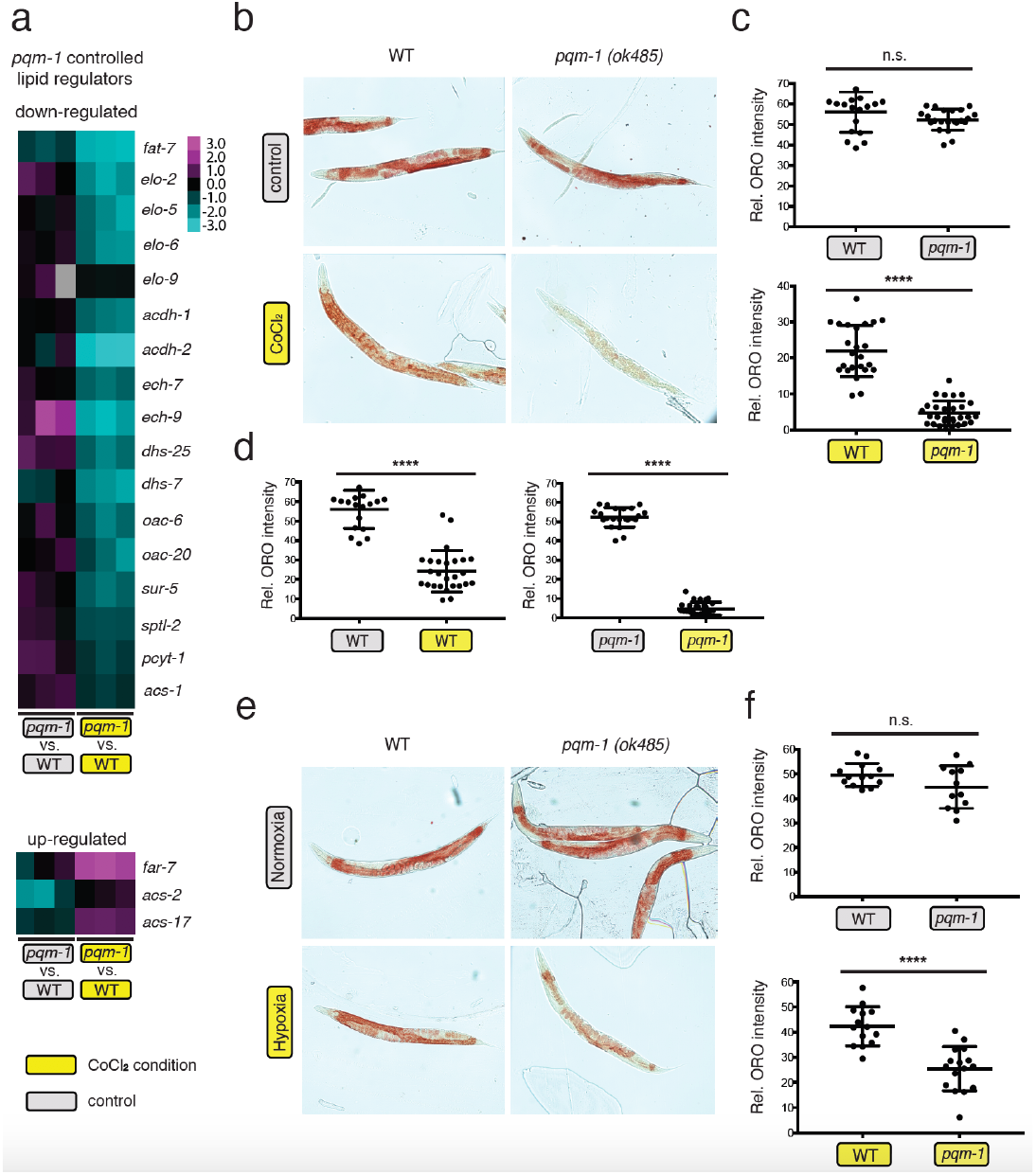
PQM-1 controls expression of lipid regulators and positively regulates hypoxic fat levels. a) Expression based analysis of *pqm-1*-dependent regulators implicated in hypoxic lipid metabolism. Two-Class SAM (*pqm-1(lf)* +CoCl_2_ versus WT +CoCl_2_ compared to *pqm-1(lf)* -CoCl_2_ versus WT -CoCl_2_) identified downregulated (top) and upregulated (bottom) lipid regulators. b-f) Oil Red O-based lipid staining of animals subjected to control conditions, hypoxia-like conditions (2.5 mM CoCl_2_, b-d), and real hypoxia (0.4% oxygen, e-f) at the early day-1 of adulthood stage for 44hr followed by Oil Red O-based lipid staining (b,e) and whole worm quantification of fat levels (c,d,f). Two-tailed *t*-test, mean ± SEM, **** p<0.0001, ns = not significant.

### The stearoyl-CoA desaturase FAT-7 functions as a downstream PQM-1 target in hypoxic lipid metabolism

The stearoyl-CoA desaturase *fat-*7 was the most significantly down-regulated lipid regulator upon exposure to the hypoxia mimetic (Fig. 4a, Fig. S4a, Table S4), suggesting that reduction of *fat-7* might be important for *pqm-1(lf)*-mediated hypoxic fat loss and subsequent survival. FAT-7 is a stearoyl-CoA desaturase that catalyzes a critical rate limiting step in fatty acid biosynthesis, desaturating stearate [CH_3_(CH2)_16_COO^−^] to mono-unsaturated oleate [CH_3_(CH_2_)_7_CH=CH(CH_2_)_7_COO^−^], which requires oxygen as an electron acceptor (Fig S4b). To verify our transcriptional results, we examined the expression of a *fat-7p::fat-7::gfp* translational reporter^24^ in *pqm-1* mutants, and found that fluorescence is significantly reduced under hypoxia-like conditions (Fig. 4b, Fig. S4c). Together, these data point to an important role of PQM-1 in *fat-7* transcriptional regulation in hypoxic conditions.

**Figure 4.**
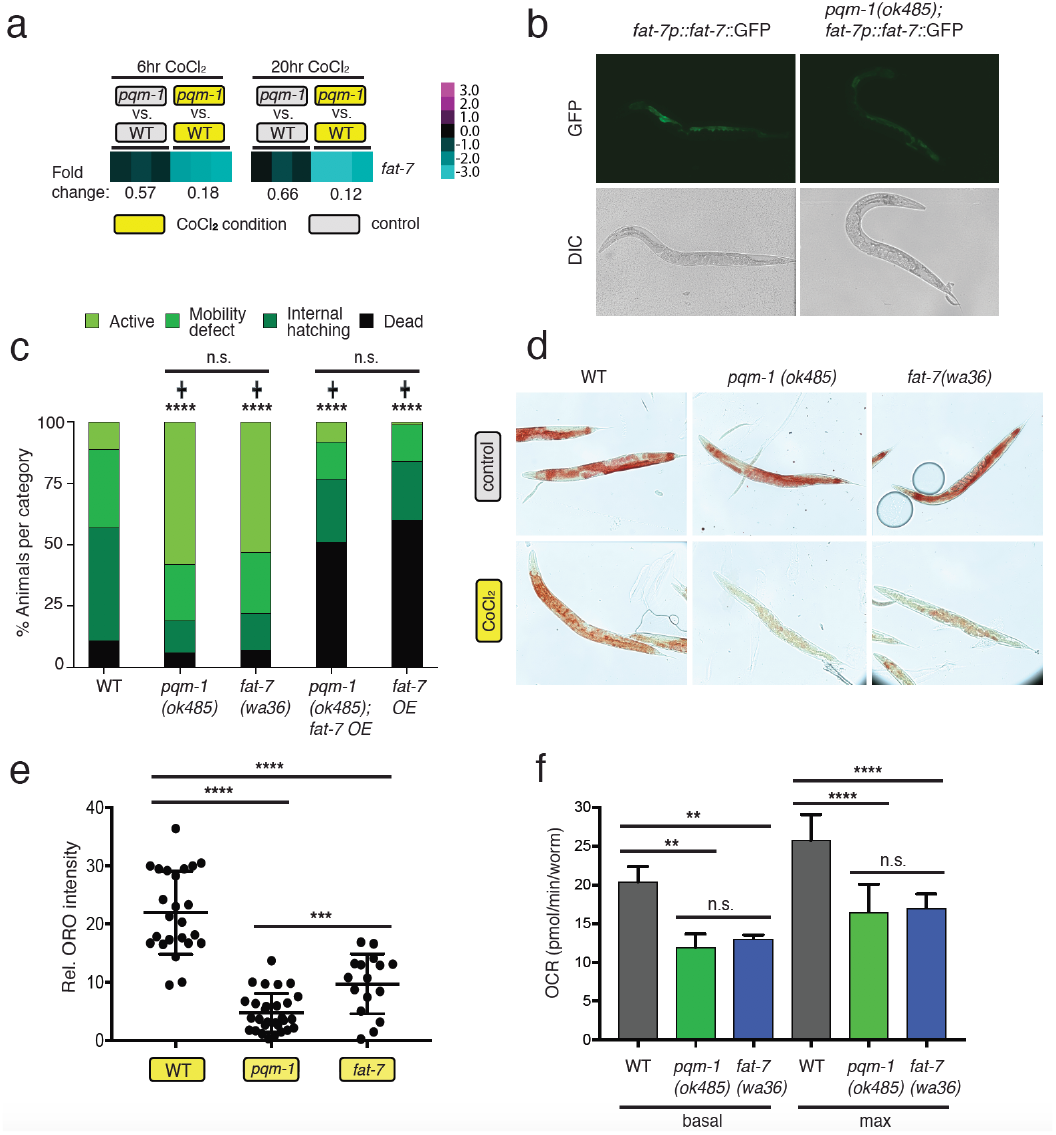
The stearoyl-CoA desaturase FAT-7 functions as a downstream PQM-1 target regulating hypoxic fat levels, oxygen consumption rates and survival. a) Gene expression analysis-based heat maps indicate a PQM-1 and CoCl_2_-dependent expression of *fat-7* upon 6 hr and 20 hr exposure to 5 mM CoCl_2_. b) PQM-1 positively regulates *fat-7* expression in hypoxia-like conditions based on a *fat-7p::fat-7::*GFP translational reporter. c) Survival and matricide analysis of indicated *C. elegans* strains exposed to 5 mM CoCl_2_ at the early adulthood stage (early day-1 of adulthood stage) for 50 hr. 61 ≤ n ≤ 134, **** p<0.0001 (**+**) indicates mutant versus wild type strain, Chi-square analysis). d) Oil Red O-based lipid staining of animals subjected to control and hypoxia-like conditions. Animals were exposed to CoCl_2_ at the early day-1 of adulthood stage for 40 hr followed by Oil Red O staining (d) and whole worm quantification of fat levels (f). g) Measurement of oxygen consumption rates. The uncoupler FCCP was injected twice (10 μM each injection) followed by sodium acid injection (NaN_3_, 40 mM, Fig. S4d). Two-tailed *t*-test, mean ± SEM, **** p<0.0001, *** p<0.001, ** p<0.01 (e,f). Three independent experiments were performed.

Our gene expression analysis suggests that *fat-7* is an essential transcriptional target of PQM-1. We tested the survival of the *fat-7(wa36)* loss-of-function mutant under CoCl_2_ conditions, and found that *fat-7* mutants displayed a moderate increase in survival when exposed to CoCl_2_ at a late larval (L4) stage (Fig. S4d). When challenged with CoCl_2_ at the early adulthood stage, which promotes matricide in wild-type worms, *fat-7* loss-of-function mutants largely phenocopied the beneficial effect of *pqm-1* ablation in hypoxia-like survival (Fig. 4c, supplementary videos 1-3). After 50 hours on CoCl_2_, approximately half of *fat-7* mutants were still actively crawling on plates, which was similar to the mobility of *pqm-1(lf)* animals under hypoxia, and strikingly different from the immobility that wild-type animals display under hypoxia-like conditions. By contrast, reintroducing *fat-7* activity into a *pqm-1* loss-of-function mutant by overexpressing a *fat-7p::fat-7::gfp* translational reporter completely reversed the beneficial effects of a *pqm-1(lf)* mutant on hypoxic mobility and survival (Fig. 4c). Overexpression of FAT-7 in wild-type and *pqm-1* worms also caused high rates of matricide and death on CoCl_2_.

Next, we asked what role *fat-7* plays in lipid regulation under hypoxic conditions. *fat-7(wa36)* mutants under normoxia did not significantly differ in fat levels relative to wild-type animals (Fig. 4d, Fig. S4e), but lack of *fat-7* activity under hypoxia-like conditions caused a reduction of lipid levels, similar to those observed in *pqm-1(lf)* mutants (Fig. 4d, e).

### *fat-7* mutants mimic *pqm-1* mutant reduction of oxygen consumption rates

We hypothesized that animals that survive in low-oxygen conditions might do so by reducing their oxygen metabolism. We measured the basal and maximum oxygen consumption rates (OCR) of wild-type and *pqm-1* worms, and found that inactivation of *pqm-1* diminished both the basal and the maximum OCR (Fig. 4f). We then found that *fat-7(wa36)* loss-of-function mutants reduced oxygen consumption to levels comparable to *pqm-1(ok485)* mutants (Fig. 4f). These data indicate that changes in FAT-7 activity result in diminished oxygen consumption in worms. Together, these data suggest that *pqm-1* mutants metabolize oxygen at lower rates due to decreased FAT-7 levels, which in turn may decrease the negative impact of low oxygen conditions on the worms. *fat-7* appears to be an important target downstream of PQM-1 in the hypoxic response and in *pqm-1’s* oxygen utilization.

### PQM-1 promotes progeny formation and survival in hypoxia-like stress

Our findings suggested that normal PQM-1 activity is detrimental for hermaphrodites exposed to hypoxia-like stress through its promotion of fat metabolism and subsequent reduction of survival. It is not immediately obvious why the normal activity of PQM-1 should be detrimental under a condition that worms might encounter in the wild with some frequency. Therefore, we investigated PQM-1’s effect on progeny survival under hypoxic conditions.

To study the effect of hypoxia on progeny development, we examined the formation of embryos in the uterus of hermaphrodites exposed to CoCl_2_. Usually *C. elegans* lays eggs that have developed to the gastrula stage, but hypoxia-like stress caused *in utero* retention of embryos that had developed further than normoxic control embryos (Fig. 5a, b). Moreover, hypoxic wild-type animals contained substantial amounts of lipid reallocating to the progeny (Fig. 5a). By contrast, hypoxic *pqm-1* mutants contained less fat, embryos in the uterus were found largely at earlier developmental stages, and the mutants displayed less internal hatching compared to wild-type hypoxic hermaphrodites (Fig. 5a, b). To examine the lipid levels in embryos, we dissected eggs out of ORO-stained hermaphrodites. Although embryos of *pqm-1* mutants had slightly reduced fat levels compared to wild-type worms under normoxic conditions (Fig. 5c,d), the effect of lipid loss was more pronounced in hypoxic *pqm-1* loss-of-function embryos. This observation suggests that PQM-1 activity normally promotes progeny development under hypoxic conditions by promoting lipid accumulation in embryos.

**Figure 5.**
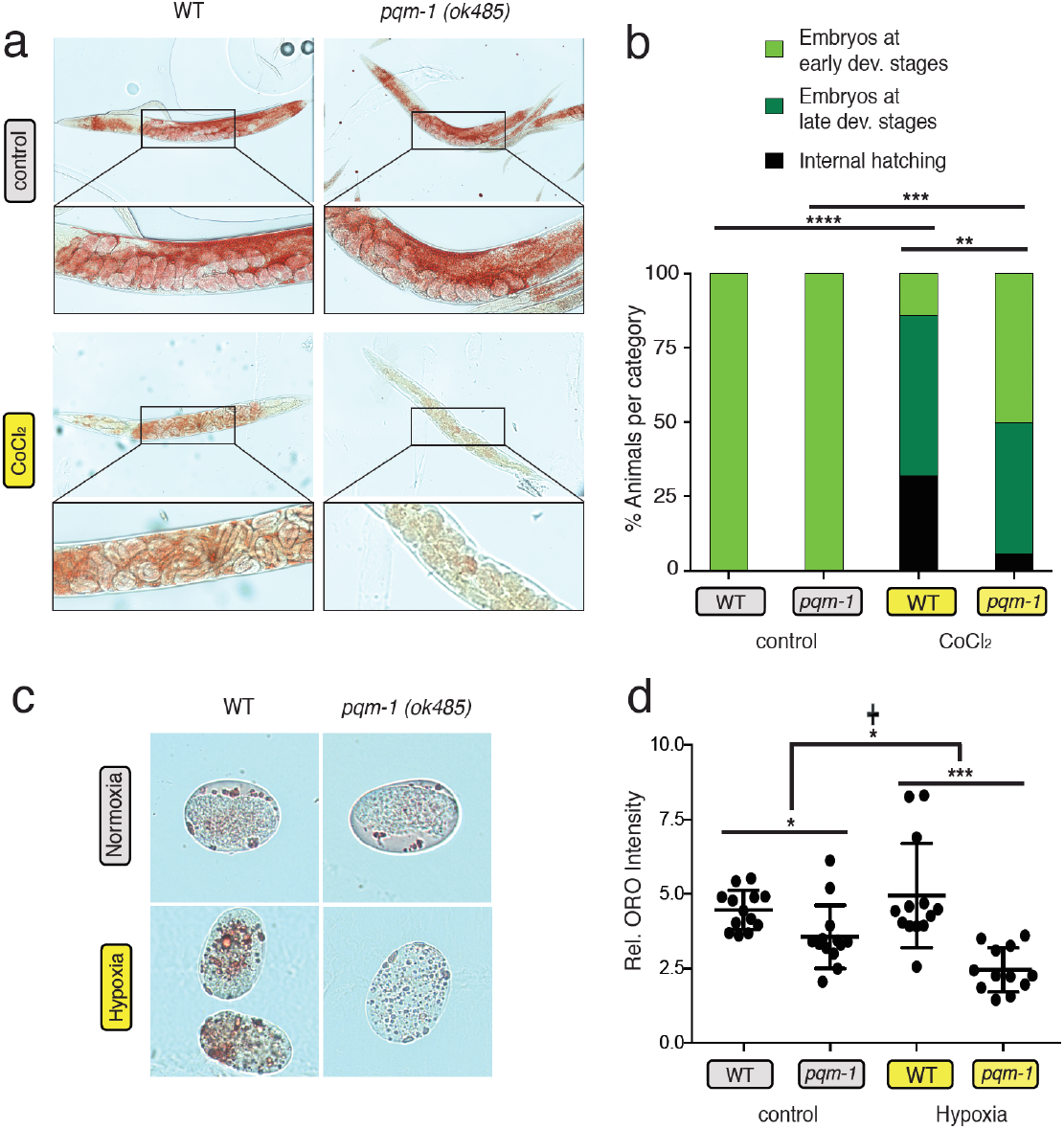
PQM-1 activity regulates progeny formation in hypoxia-like stress. a-b) Oil Red O-based lipid staining displaying progeny in parental hermaphrodites (a) and quantification of matricide and embryonal stages in the uterus of hermaphrodites (b). **** p<0.0001, *** p<0.001, ** p<0.01, Chi-square analysis was performed. Parental hermaphrodites were Oil Red O strained at the day-1 of adulthood stage (control). CoCl_2_ challenged hermaphrodites were exposed to 2.5 mM CoCl_2_ at the day-1 of adulthood stage for 44 hr followed by Oil Red O staining. c-d) Lipid content of eggs (c) dissected from Oil Red O stained hermaphrodites challenged for 44 hr with 0.4% oxygen and quantification of fat levels in eggs (d). Two-tailed *t*-test, mean ± SEM, * p<0.05, *** p<0.0001. Two-way ANOVA (**+**), p=0.02.

Internal hatching of progeny in *C. elegans,* also known as “matricide,” has been proposed to be an adaptive response to stress or starvation^25^, enabling the parent to provide nutrients for larval development^26^ at the cost of the mother’s life. To assess the role of matricide in hypoxic survival, we determined the fraction of worms displaying internal hatching during survival assays on CoCl_2_. *pqm-1* inactivation reduced the occurrence of matricide, and inhibited the high matricide rate of *daf-16(lf)* mutants (Fig. 6a). Similarly, inactivation of *fat-7* reduced internal hatching under hypoxia-like stress (Fig. 6b), suggesting that the conversion of saturated fatty acids (SFAs) to mono-unsaturated fatty acids (MUFAs) is required for promoting matricide. Interestingly, the total lipid level and the occurrence of matricide are positively correlated in hypoxic wild-type worms, *pqm-1(lf),* and *fat-7(lf)* mutants (Fig. 6c), suggesting an important role of fat metabolism in internal hatching and progeny development.

**Figure 6.**
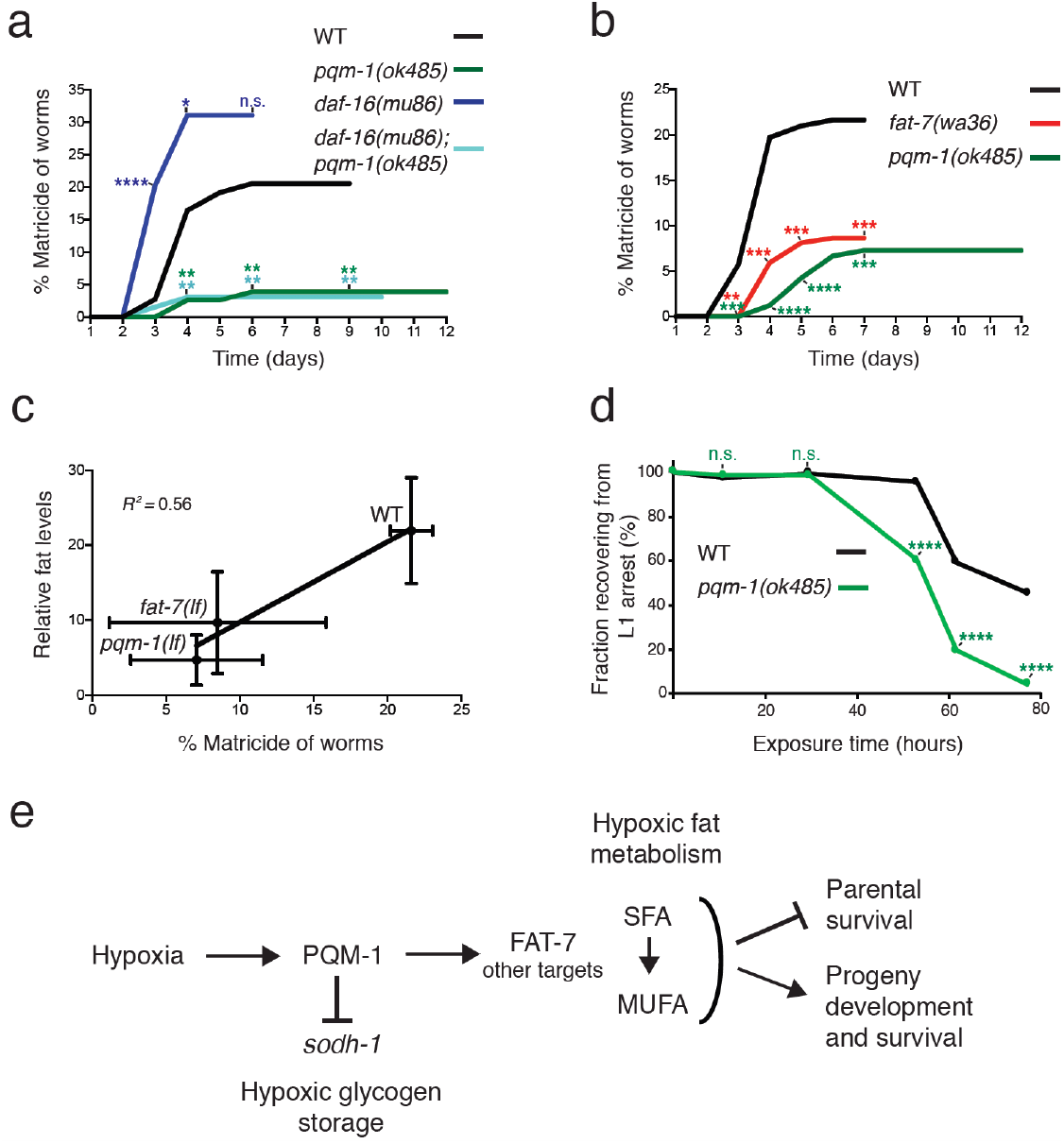
PQM-1 activity promotes matricide of hermaphrodites and progeny survival in hypoxia-like stress. a, b) Matricide analysis of C. elegans strains exposed to 5 mM CoCl_2_ at the L4 larval stage. 65 ≤ n ≤ 90 (a), 157 ≤ n ≤ 185 (b). At least two independent experiments were performed. c) Matricide positively correlates with fat levels of indicated strains. d) Survival analysis of progeny originating from CoCl_2_ exposed hermaphrodites. The ability to exit and survive a developmental arrest caused by CoCl_2_ was analyzed after transferring arrested larvae to regular culture conditions. **** p<0.0001, *** p<0.001, ** p<0.01, * p<0.05 indicates mutant versus wild type strain, ns = not significant. Chi-square analysis. Three independent experiments were performed. e) Model for PQM-1 as a regulator of hypoxic lipid and glycogen metabolism and survival. PQM-1 activity limits parental survival through promoting lipid metabolism, whereas it supports progeny formation and survival in hypoxialike conditions. PQM-1 activity represses sorbitol dehydrogenase (sodh-1) expression affecting hypoxic glycogen storage. SFA, saturated fatty acid: stearic acid [CH_3_(CH2)_16_COOH]; MUFA, mono-unsaturated fatty acid: oleic acid [CH_3_(CH_2_)_7_CH=CH(CH_2_)_7_COOH].

To address whether PQM-1 is also required for progeny function under hypoxia, we analyzed the ability of progeny to survive and recover from hypoxia-like stress. Wild-type *C. elegans* larvae hatched from eggs that are laid by CoCl_2_-exposed hermaphrodites arrest as L1s if maintained on CoCl_2_. These L1-arrested progeny are able to recover and develop into reproductive adults when they are transferred to regular culture conditions (Fig. 6d). However, *pqm-1* progeny of CoCl_2_-treated mothers are impaired in their ability to exit L1 larval arrest (Fig. 6d), indicating that PQM-1 activity is essential for prolonged progeny survival under hypoxic stress and subsequent recovery. Taken together, our results point to a balancing role for PQM-1 in hypoxia-like conditions, promoting progeny survival at the cost of somatic integrity of the parental generation (Fig. 6e).

## Discussion

Here we have found that the zinc finger transcription factor PQM-1 is a regulator of lipid metabolism and subsequent survival in hypoxic conditions, affecting both parental and progeny survival. Organisms obtain energy by metabolizing energy sources through the process of respiration, and fatty acids and carbohydrate are the major energetic substrates for ATP production. The volume of oxygen consumed, and the volume of carbon dioxide produced (the “respiratory quotient”) depends on the fuel source. Molecules that are less oxidized, such as fatty acids, require more oxygen to be metabolized to CO_2_ and H_2_O than do fuel sources that are more oxidized, such as carbohydrates. Although lipid oxidation provides more ATP than carbohydrates, it also requires more oxygen per mole of ATP synthesized^27^. Therefore, glycolysis, especially anaerobic glycolysis, is an oxygen-saving but less effective process for energy production, in contrast to lipid catabolism, which consumes oxygen through oxidation of lipids in mitochondria^27^. When oxygen is abundant, fat metabolism is highly efficient and is thus the preferred catabolic pathway; but in hypoxic conditions, the relatively high rate of oxygen consumption can have deleterious effects on a tissue or an organism. Under hypoxic conditions, however, glycogen is the primary energy source for *C. elegans* in anoxia^17^. We found that *pqm-1(lf)* mutants maintain higher glycogen levels in hypoxia-like conditions. Inactivation of sorbitol dehydrogenase-1 (*sodh-1*), which is strongly upregulated in the absence of *pqm-1,* diminished the elevated glycogen content of a *pqm-1(lf)* mutant. SODH-1 enzymatic activity converts sorbitol, the sugar alcohol from of glucose, into fructose^19^. In previous studies it has been demonstrated that sorbitol can be metabolized to glycogen in diapause eggs of insects (*Bombyx mori*) depending on the state of the diapause^23^. SODH-1 activity is upregulated at the termination of the diapause and metabolizes sorbitol to fructose, which is further converted to glycogen^23,28,29^. We anticipate that a similar sorbitol-to-glycogen metabolic pathway functions in *C. elegans.* Consistency, our data indicate that loss of *pqm-1* and thus elevated *sodh-1* expression is associated with increased glycogen storage, while inactivation of *pqm-1* resulted in a reduction of *fat-7* desaturase expression and diminished lipid levels under hypoxic stress.

The enzymatic reaction that FAT-7 carries out, desaturation of stearic acid to oleic acid, requires molecular oxygen^30^, and our data suggest that basal and maximal oxygen consumption rates are decreased in both *pqm-1* and *fat-7* mutants. A functional link between the desaturation of lipids and respiration has been previously established in plants^31^, through a deficiency in an ω-6-oleate desaturase, which regulates lipid metabolism and respiration. Similarly, we found that basal and maximal oxygen consumption rates are decreased in both *pqm-1* and *fat-7* mutants. Our data suggest that PQM-1 might act as a metabolic gatekeeper in hypoxia, repressing *sodh-1* expression and glycogen levels, while promoting the expression of the stearoyl-CoA desaturase FAT-7. FAT-7 activity, in turn, regulates lipid levels and their desaturation, which appears to be required for respiratory activity and oxygen consumption.

The embryonic heart is a prominent example of suppression of lipid metabolism in a hypoxic tissue that largely relies on carbohydrates (glucose) as an energy source^32^. The developing heart of an embryo is exposed to hypoxia; however, it is highly protected against hypoxic stress through the reduction of lipid metabolic processes^33^. Downregulation of fat metabolism by the activity of the basic helix-loop-helix transcription factor HAND1 decreases both basal and maximal OCRs. A lower oxygen consumption rate in cardiomyocytes provides a protective metabolic strategy for tissue development under low oxygen tension. Inhibition of cellular lipid metabolism by etomoxir, a carnitine palmitoyltransferase antagonist that prevents mitochondrial long-chain fatty acid import, also decreases OCR and protects against myocardial ischemia^34,35^; downregulation of lipid metabolism and OCRs resulted in a shift of energy metabolism to glycolysis.

Why has a PQM-1-mediated mechanism evolved, which boosts hypoxic fat levels if this seems to have negative implications for survival under hypoxic stress? We hypothesize that PQM-1 positively regulates intestinal fat production to fuel reproductive processes in hypoxic animals. An investment in reproduction is beneficial for the population, even if detrimental to the individual mother. Such a strategy might enable some progeny to potentially escape hypoxic stress conditions, thereby ensuring survival of the population. A similar trade-off between viability and fecundity has been previously described when *C. elegans* is exposed to nutrient-poor and oxidative stress environments^26^. During reproduction, somatic resources, particularly lipids, are reallocated to the germline. Fat is produced in the intestine and the hypodermis and transported by the actions of vitellogenins^36^, which assemble with transport lipids in the form of yolk to shuttle fat from the intestine to the developing oocytes. When resources are limited, lipid reallocation appears to promote fecundity at the cost of somatic integrity of the parental generation. We found that both vitellogenins were downregulated in *pqm-1* mutants upon hypoxic stress (Table S2); thus, vitellogenin genes are normally positively regulated by PQM-1 upon hypoxic stress, perhaps in order to facilitate lipid transport from the mother’s intestine into developing eggs. Previously, it has been demonstrated that PQM-1 suppresses vitellogenin expression and acts as a downstream transcriptional effector of TORC-2 signaling^37^. However, these studies were performed under regular growth condition for *C. elegans*, and not under hypoxic stress. Thus, it is possible that PQM-1 controls vitellogenin expression depending on environmental impacts and responds in differently under stress versus standard growth conditions for worms. Taken together, we found that PQM-1 activity increases hypoxic lipid levels in worms by positively regulating *fat-7* expression and promotes the reallocation of fat to embryos under hypoxic stress most likely via controlling vitellogenin expression. This, in turn, boosts lipid-dependent embryonic development inside the worm’s uterus (Fig. 5a, b; 6a, b). Somatic allocation of limited lipid resources into the promotion of reproduction during hypoxic stress is detrimental for individual viability, but may increase the chance of species survival through investment in the next generation.

To address the effect of hypoxia on PQM-1’s metabolic readout, we have used chronic (lower level) hypoxia and the hypoxia mimetic CoCl_2_. CoCl_2_ activates HIF-1^16^, which might mimic HIF-1-dependent chronically lower levels of hypoxia, rather than severe anoxic conditions, which trigger HIF-1-independent survival responses in *C. elegans*^7^. Several solid tumor types in humans generate a microenvironment inside the tumor where cancer cells are exposed to lower levels of hypoxia (approximately 1% atmospheric oxygen)^38^. We anticipate that CoCl_2_ might mimic such a HIF-dependent, persistent hypoxia response relevant for cancer cell metabolism. Because lipid and carbohydrate-related pathways are well conserved across species, deciphering mechanisms implicated metabolic remodeling in *C. elegans* will be beneficial for the development of treatment strategies to treat cancer and age-related diseases in humans.

## Materials and Methods

### *C. elegans* genetics

All strains were cultured using standard methods^39^. In all experiments, N2 is wild type. LG II: *pqm-1(ok485).* LG III: *daf-2(e1370). daf-2 RNAi* was performed in an OP50 bacterial background^40^.

### Strains

CQ200 (*pqm-1(ok485*);*daf-2(e1370)*); CQ528 (*pqm-1 (ok485)*); CF1041 (*daf-2(e1370*)); CQ565 (*daf-16(mu86);pqm-1(ok485)*); DMS303 (*nIs590 [Pfat-7::fat-7::GFP* + *lin-15(*+*)]); RB8615 (pqm-1(ok485*);(*nIs590 [Pfat-7::fat-7::GFP* + *lin-15(*+)]); BX153 (*fat-7(wa36)*); RB8610 (*sodh-1(ok2799)*); RB8611 (*pqm-1(ok485);sodh-1(ok2799)*); CF1038 (*daf-16 (mu86)*); CF2124 (*muIs139 [dod-11p::RFP(NLS) + rol-6(su1006)]*); All animals were synchronized at the L4 larval stage for adult analyses, as *pqm-1* mutants develop more slowly than do wild-type animals^15^.

### Survival analysis

Synchronized L4s were picked onto plates to eliminate any contribution of delayed development into adult survival analysis. The first day of adulthood was defined as t=0, and the log-rank (Mantel-Cox) method was used to test the null hypothesis in Kaplan-Meier survival analysis, as previously described ^41^, and evaluated using JMP survival analysis software ^42^. The log cumulative hazard function was also estimated. All experiments were carried out at 20°C; n ≥ 60 per strain/trial.

### Cobalt chloride-based survival assays

A 200 mM cobalt chloride (CoCl_2_) stock solution was prepared, and filter sterilized using a 0.22 μm filter as described previously^8^. OP50 bacteria (60 μl bacteria solution) were seeded in the center of 6 cm NG plates previously dried for 3 to 4 days at room temperature. OP50 bacteria were grown on plates overnight at room temperature. To prepare CoCl_2_ plates, 250 μl of the 200 mM CoCl_2_ stock solution was added to 6 cm NG plates (contain ^~^10 ml NG agar) with seeded, over-night grown OP50 bacteria, which results in a 5 mM final concentration of CoCl_2_. The CoCl_2_ solution was immediately distributed equally over the surface of the plate with a spreader (carefully, not to remove seeded OP50). The CoCl_2_ solution was completely taken up by the plate resulting in dry plates before worms were added. A thin ring of 100 % glycerol was added along the plastic wall of the plate to reduce the number of worms crawling off the agar. Plates were incubated with lids facing up, which can reduce the number of worms crawling off CoCl_2_ containing plates. To obtain synchronized worm cultures, 60 to 100 L4 larvae were picked for each strain, grown for 4 hours on OP50 bacteria and added to freshly prepared CoCl_2_ plates (at a 4 hours-post-L4 larvae stage, corresponds to first day in survival assay). The log-rank (Mantel-Cox) method was used to test the null hypothesis in Kaplan-Meier survival analysis, as previously described ^41^, and evaluated using JMP survival analysis software ^42^. The log cumulative hazard function was also estimated. Internally hatched worms were censored for survival assays.

### Cobalt chloride-based Matricide Assays

L4 synchronized worms and CoCl_2_ containing plates were prepared as described above for survival assays. Worms are exposed to CoCl_2_ at the L4 stage (Fig. 6a,b) or the day 1 of adulthood stage (Fig. 4c). Exposure of worms at the L4 stage results in reduced matricide rates over time (e.g. ^~^20-25% for wild type worms), enabling the calculation of matricide curves (Fig. 6a,b), whereas challenging day 1 adult worms with CoCl_2_ causes already higher rates of internal hatching after 2 days of CoCl_2_ exposure (Fig. 4c).

### Oil Red O-based lipid staining and quantification

L4 synchronized worms were grown to day 1 of adulthood. OP50 Plates were prepared as described above. CoCl_2_ solution was added to plates (2.5 mM final concentration). 1 hour after adding CoCl_2_ to plates seeded with OP50 worms were transferred to CoCl_2_ plates and exposed to CoCl_2_ conditions for 40 hr. A 0.5% Oil Red O stock solution was prepared in high-quality 100 % isopropanol as described previously^43^. For staining of worms, the stock solution was diluted to 60 % with sterile water, incubated on a rocking platform at room temperature overnight and filtered through a 0.45 μm filter. Worms were resuspended in 500 μl 60 % isopropanol for fixation. Isopropanol was aspirated, 500 μl of freshly filtered Oil Red O working solution was added and worm strains were incubated in a Thermomixer at 25°C using mild agitation (550 rpm) for 16 hours. Animals were washed 3 times with 500 μl of 0.01 % Triton X-100 in M9 buffer and stored at 4°C followed by imaging on a Nikon Eclipse Ti microscope (10x objective). Oil Red O quantification was performed as previously described^44^. In brief, color images were split into RGB monochromatic images in Image J. The Oil Red intensity was determined by calculating the mean gray value within a worm region (Intensity of the blue channel was used as the signal, adjusted by the intensity in the Red channel as the background).

### RNA collection and Microarray Hybridization and Analysis

Worms were collected, RNA isolated and hybridized on 4×44K *C. elegans* arrays (Agilent) at 60°C overnight, as previously described^45^. Three biological replicates of *pqm-1(ok435)* on CoCl_2_ versus wild type on CoCl_2_ were compared to *pqm-1(ok435)* without CoCl_2_ versus wild type without CoCl_2_. In addition, *pqm-1(ok435)* on CoCl_2_ versus *pqm-1(ok435)* without CoCl_2_ were compared to WT on CoCl_2_ versus WT without CoCl_2_. Significant differentially-expressed gene sets were identified using one or two-class SAM^46^. Enriched motifs were found using RSAT^47^. Gene ontology analysis (GO) was performed utilizing DAVID (Database for Annotation, Visualization and Integrated Discovery)^48^ and g:Profiler^49^. GO categories were visualized with REVIGO (Reduce and Visualize Gene Ontology),^50^. Microarray data can be found in PUMAdb (http://puma.princeton.edu).

### Glycogen staining

Glycogen staining was performed as previously described^17^. Worm strains were exposed at the L4 larval stage to 5 mM CoCl_2_ for 3 days. Strains were pairwise compared by picking each of them into an M9 droplet on an agar pad (two small agar pads were placed next to each other on the same slide). As soon as the M9 droplets were largely evaporated, the pad was inverted over the opening of a 50 g bottle of iodine crystal chips (Sigma, St. Louis, MO) for 1 min. After the iodine stained color of the agar pad disappeared (non-specific staining), worms (about 10-15 worms per treatment) were immediately imaged by a Nikon Eclipse Ti microscope.

### Measurement of oxygen consumption rate (OCR)

OCR measurement was carried out as previously described^51^. Worms were synchronized at the L4 stage and OCR measurement was performed at early adulthood (day 1 of adulthood stage). Approximately 15-20 animals were pipetted into each well of a Seahorse XF96 utility plate. 6 replicates per strain were used (6 technical repeats). The Seahorse program was set up in a way that each oxygen consumption measurement consisted of a two-minute mix cycle, followed by a one-minute wait period (to allow worms to settle), and finally a two-minute interval for measurement of oxygen levels. OCRs were normalized by worm number and worm body area. To determine the basal OCR we averaged the first tree measurements (Fig. S4d). The uncoupler FCCP was injected twice (10 μM each injection) followed by sodium acid injection (40 mM). The first two measurements after FCCP injection were variable. Therefore, we averaged measurements 3-10 after FCCP injection to obtain the maximal OCR.

### Hypoxic Incubation

Survival analysis of worms in a hypoxic glove box (Coy Laboratory Products) were performed as previously published^52^. Worms were exposed on NG plates to 0.3 % O_2_ (balanced with nitrogen) for 25 hours at 26°C followed by a one-day recovery period in room air at 20°C.

### L1 arrest-based survival assay of progeny

L1 larvae hatched from eggs laid on OP50 plates containing 5 mM CoCl_2_. Larvae did not develop further and arrested when exposed to CoCl_2_. 60 to 120 arrested larvae per strain were transferred every day to fresh OP50 plates without CoCl_2_ and analyzed for their ability to exit and survive the larval arrest and develop to adulthood.

### RNAi treatment

RNAi-treated worm strains were fed *E. coli* (OP50)^40^ containing an empty vector construct or a construct expressing double-stranded RNA (dsRNA) against the gene of interest.

## Acknowledgments

This work was supported by the NIH NIA 1R56AG047344-01A1 293 (C.T.M.) and the Glenn Foundation for Medical Research. C.T.M. is the Director of the Glenn Center for Aging Research at Princeton, which also supported T.H. We thank the *Caenorhabditis* Genetics center for providing *C. elegans* strains, and Jasmine Ashraf for technical assistance. We also thank members of the C.T.M. laboratory for comments on the manuscript.

## Author Contribution

T.H. and C.T.M. designed experiments. T.H. performed experiments. T.H. and C.T.M. wrote the manuscript.

## Notes

http://puma.princeton.edu

