## Supplemental Figures 1-5 for "PQM-1 controls hypoxic survival via regulation of lipid metabolism"

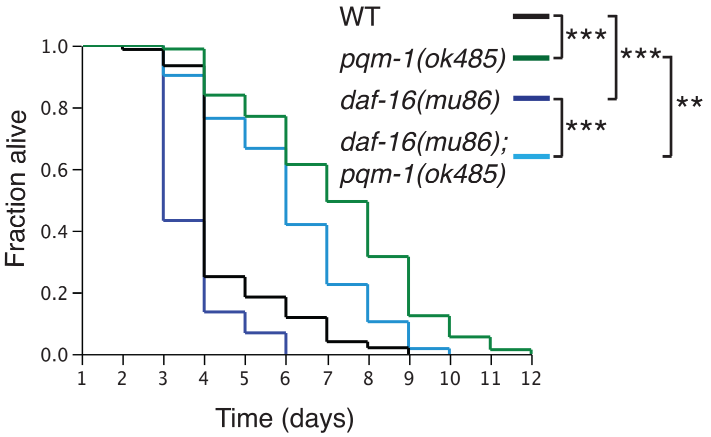


Supplemental Figure 1

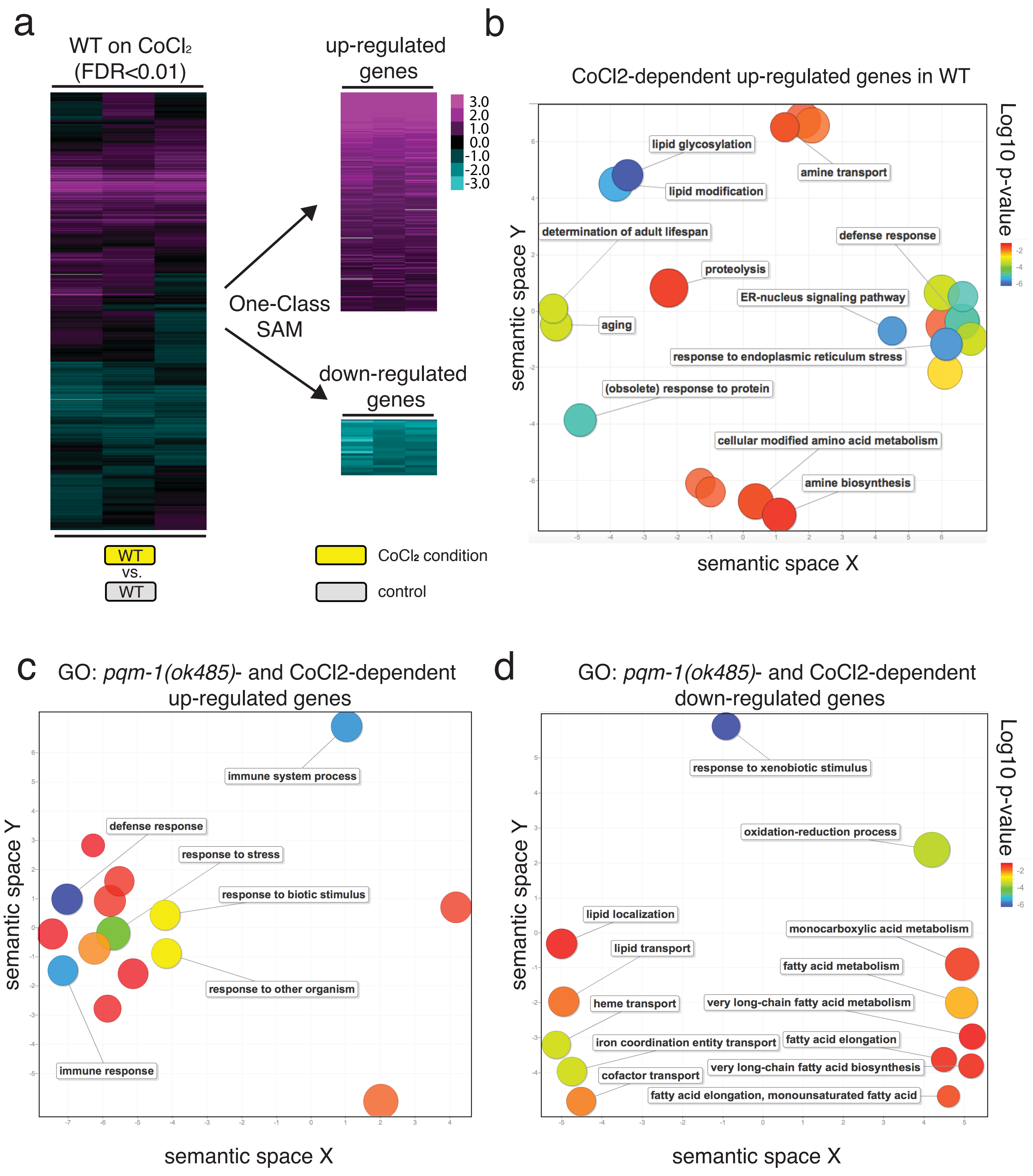


Supplemental Figure 2

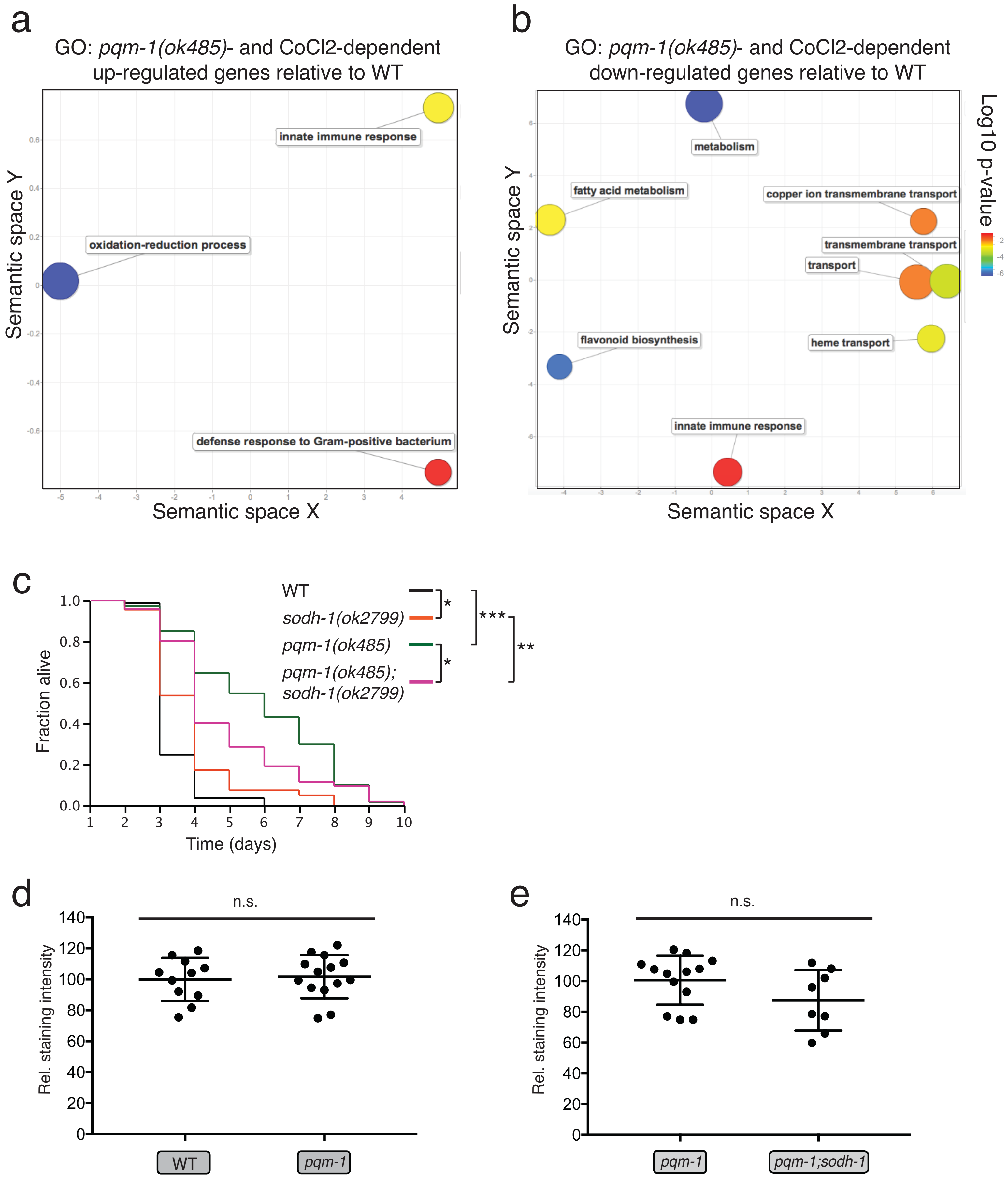


Supplemental Figure 3

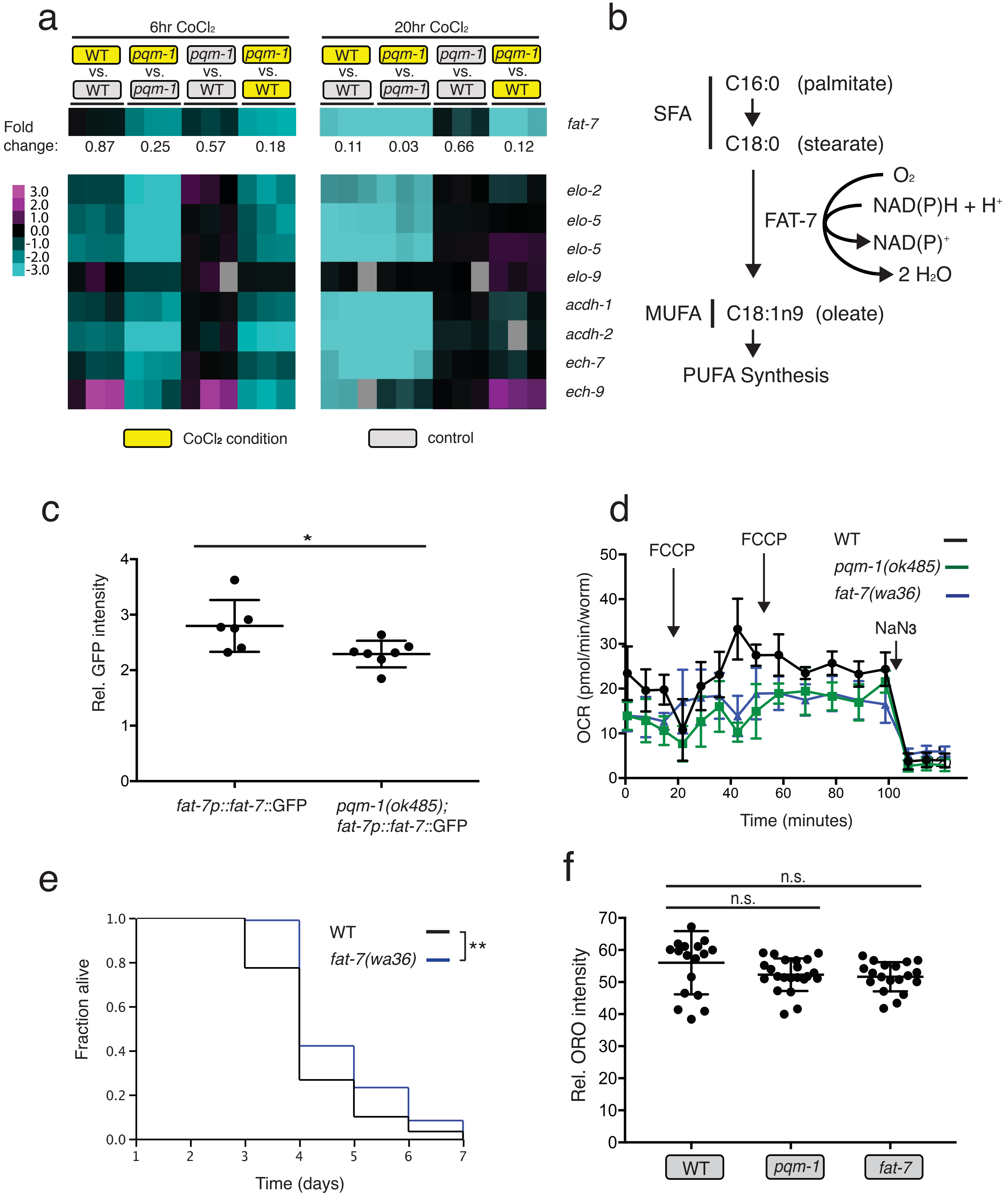


Supplemental Figure 4

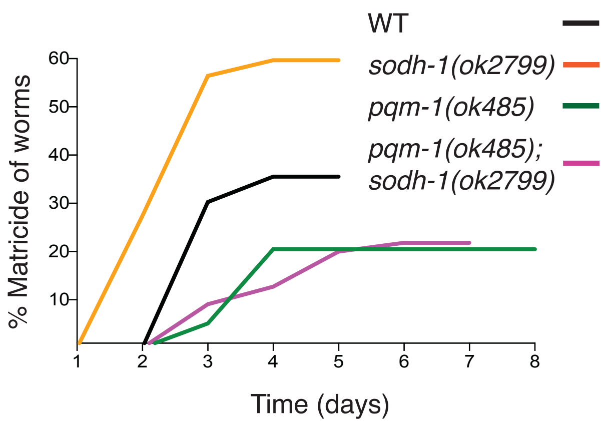


Supplemental Figure 5
